# EEG covariance patterns detects global cortical activity distribution following weak tactile inputs

**DOI:** 10.1101/2023.11.06.565763

**Authors:** Astrid Mellbin, Udaya Rongala, Henrik Jörntell, Fredrik Bengtsson

## Abstract

Many studies have suggested that the neocortex operates as a global network of functionally interconnected neurons, indicating that any sensory input could shift activity distributions across the whole brain. A tool assessing the activity distribution across cortical regions with high temporal resolution could then potentially detect subtle changes that may pass unnoticed in regionalized analyses. We used eight-channel, distributed electrocorticogram (ECoG) recordings to analyze changes in global activity distribution caused by single pulse electrical stimulations of the paw. We analyzed the temporally evolving covariance patterns of the activity distributions using principal component analysis (PCA). We found that localized tactile stimulation caused clearly measurable changes in global ECoG activity distribution. These changes in signal covariance patterns were detectable across a small number of ECoG channels, even when not including the somatosensory cortex, suggesting that the method has high sensitivity, potentially making it applicable to human EEG for detection of pathological changes.

## Introduction

Historically, the brain has been perceived as an organ with specific functions localized in distinct regions ^1^. Thus, although along somewhat different schemes, the cerebral cortex is divided into discrete areas that process different sensory, motor and cognitive in-and outputs ^2,3^. In turn these areas have been divided into areas that process primary inputs, secondary areas that integrate various aspects of information and then into associative areas that further integrate information. In the cortex, the input is thought to be processed in an orderly fashion in vertical columns ^4^ where primary sensory information is received in one part and then distributed to other parts where it is integrated with other information.

Nevertheless, alternative historical theories have posited that information processing might occur in a more distributed manner ^5,6^. Recent studies have supported this notion, revealing that the neuronal decoding of tactile sensory information extends beyond the corresponding neocortical region and is distributed across several different cortical areas ^7,8^. Information from visual input has also been found across the cortex in awake mice engaging in visual discrimination tasks ^9^. Likewise, recent research using wide-field calcium imaging has uncovered similar results, demonstrating the dispersion of cortical activity in motor control, learning, and visual processing ^10^. Furthermore, various aspects of tactile inputs can be detected in numerous parts of the thalamus ^11^.

In light of these findings, changes in activity evoked by tactile inputs should theoretically be observable across various cortical areas simultaneously, also when employing less invasive techniques. Considering the cortex as a globally interconnected network, each instance of sensory input can be likened to an injection of activity via thalamocortical synapses. This injection initiates perturbations in the current distribution of global cortical activity, potentially propagating throughout the entire cortical network. Since the distribution of activity within the cortex spontaneously evolves at a rapid pace, the resulting pattern of activation among the cortical neuron population depends not only on the spatiotemporal structure of the sensory input but also on the prevailing distribution of cortical neuron activity at the time of stimulation. Hence, the exact paths of propagation of the sensory-evoked perturbation through the global cortical neuron population ^8,10^ will depend on activity-or state-dependent network branching patterns ^12,13^.

Despite the potentially astronomical number of permutations of these branching patterns (given, for instance, that there are tens of millions of cortical neurons in rats ^12^), statistically predominant covariance patterns may still exist among these neurons, representing normal, unperturbed cortical states. In contrast, less common covariance patterns may signify perturbed states, such as those induced by sensory input or neurological diseases that disrupt these normal branching patterns of activity propagation. The extent to which such perturbed global covariance patterns can be detected via non-invasive methods is of great importance, as non-invasive techniques are widely applicable to humans and hold significant clinical potential, particularly in monitoring conditions like epilepsy, Parkinsonism, and psychosis.

Non-invasive methods have a much lower resolution than many invasive methods, i.e., they provide a subsampling of the underlying phenomenon, which in this case is the dynamics in the activity distribution across the local neuron population. Despite this, we hypothesize that if the response to an input or a perturbation is distributed globally, then it is potentially possible to use that fact as a means to extract high precision information, if the recorded signal has a high temporal resolution. Specifically, in multi-channel EEG recordings there is information about global activity distribution changes from many different locations that could potentially be used as a proxy for the underlying covariance patterns in the neuron population.

This leads to the question of how sensitive this non-invasive approach can be. Can it detect the global effects of ultra-brief sensory inputs, even when activity in the primary sensory area associated with that input type is not considered? We address this question by employing multi-channel surface ECoG recordings, which are macro-electrode recordings closely related to EEG recordings that are non-invasively applicable to humans.

## Materials and method

### Preparation

Adult male Sprague Dawley rats (n=27, weighing 324-508 grams) were initially sedated using isoflurane (3% mixed with air for 60–120 s). and anesthetized with a ketamine/xylazine mixture injected intraperitoneally (ketamine: xylazine concentration ratio of 15:1, initial dose approximately 60 mg/kg ketamine) and underwent preparatory surgery. Anesthesia was then maintained with a continuous intravenous infusion of a ketamine/xylazine mixture (approximately 5 mg/kg per hour for the ketamine component, end concentration ratio of 20:1 for ketamine: xylazine). The absence of withdrawal reflexes in response to noxious pinch of the hind paw was used to ascertain adequate levels of anesthesia until the brain was exposed. Once the brain was exposed the occurrence of sleep spindles in the ECoG was used together with continuous testing of the withdrawal reflexes to monitor that the anesthesia was maintained at an adequate level.

Four craniotomies, 5×5 mm each, were performed in order to get access to the recording areas. Two of the craniotomies were made over the sutura coronaria, bilaterally, each craniotomy providing access to the primary motor cortex and the primary sensory cortex (S1). Two craniotomies were made rostrally to the sutura lambdoidea, bilaterally, each craniotomy providing access to the primary visual cortex and the primary auditory cortex (Fig 1A). A pool of cotton in agar was built and then filled with 37℃ paraffin oil to keep the exposed parts of the brain from drying. The dura mater was cut in the rostral part of the exposed areas so that the cerebrospinal fluid (CSF) could escape, and the dura lay flat on the cortical surface. Cotton wool drains were placed over the edge of the pool so that the CSF was continuously drained from the pool.

**Figure 1.**
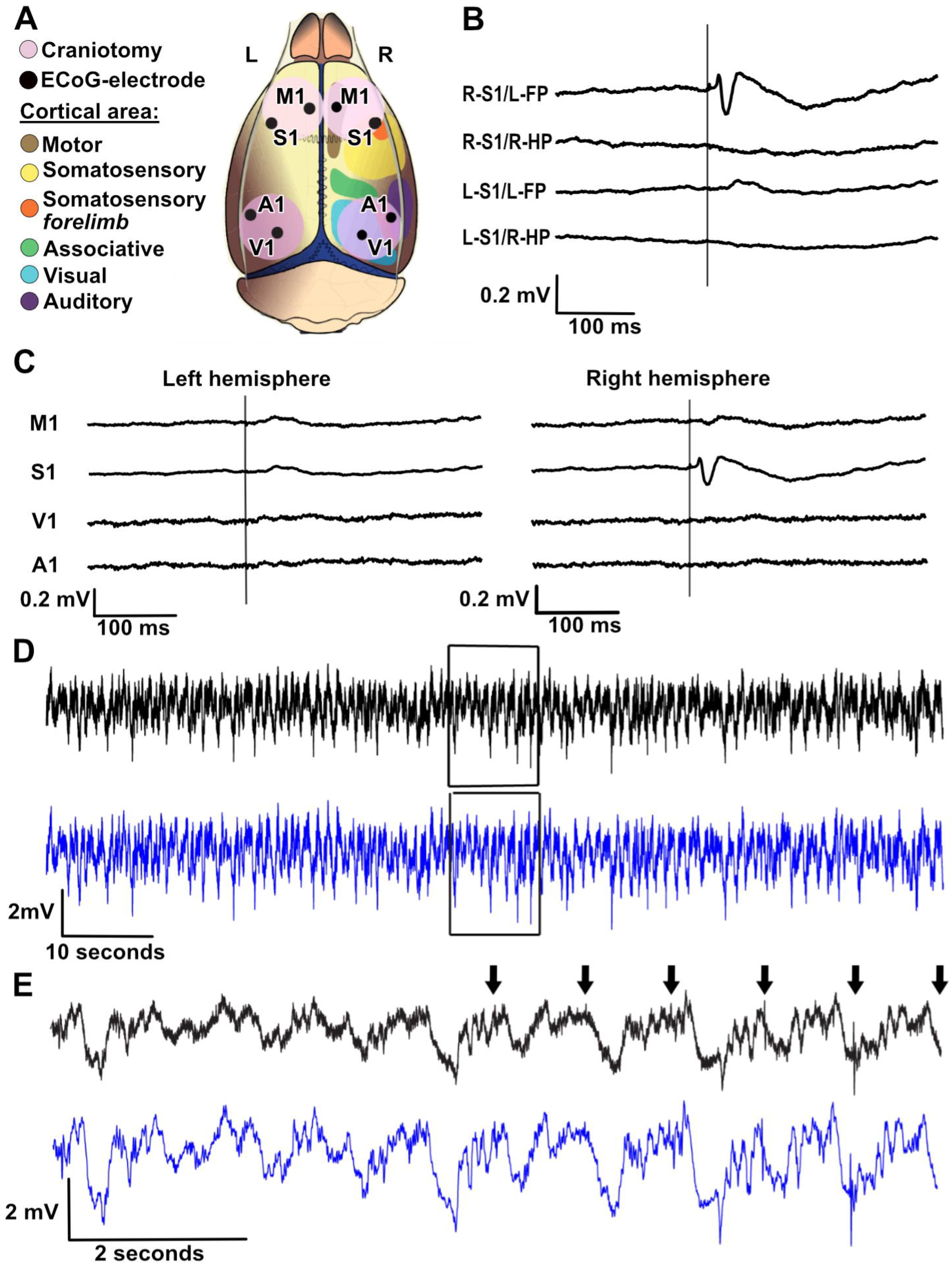
ECoG recording setup. **A** Locations of the ECoG recording electrodes relative to the outlines of the approximate location of different neocortical areas. **B** Averaged evoked cortical response (ECoG) recorded from the right and the left S1 areas. The vertical line indicates the time of stimulation. **R-S1/L-FP** is the average response evoked by electrical skin stimulation (1 Hz, 300 repetitions) of the left forepaw in the right S1. **R-S1/R-HP** is a recording from the same S1 location but with electrical skin stimulation (1 Hz, 300 reps) of the right hind paw. **L-S1/L-FP** is a recording from the left S1 during electrical skin stimulation (1 Hz, 300 reps) of the left forepaw. **L-S1/R-HP** is a recording from the same location in the left S1 with electrical skin stimulation (1 Hz, 300 reps) of the right hind paw. Note that neither of the stimulations (fore-/hind paw) evoked a response in the left S1, since both S1 recording electrodes were located in the forelimb areas of the respective hemispheres. **C** Averaged responses from the eight ECoG electrodes to stimulation of the left forepaw. **D** Long, time continuous raw ECoG trace from the right S1 (same recording point as in **B**, top two traces). Top, unfiltered recording trace. Bottom, the same trace after removal of the shock artifact and application of a moving average filter (see methods for details). **E** Zoom-in on the traces in **D (**box). The arrows indicate stimulation times for the forepaw stimulation. Top, unfiltered record trace. Bottom, after removal of the shock artifact and application of a moving average filter (see methods for details).

Eight ECoG electrodes (surface silver ball electrodes, see below) were placed in the exposed cortical areas (Fig 1A) and two grounding electrodes were placed in the neck muscles. The two S1 ECoG electrodes were placed in the forepaw areas, as evaluated by electrical stimulation of the skin of the left forepaw or right hind paw eliciting maximal field potential responses in these areas, or not (Fig 1B-C).

### Recordings

Recordings were made with silver ball electrodes (Ø 250 um). The signal was passed through a Digitimer NL844 pre-amplifier with a low frequency cut off at 0.1Hz and gain x1000, connected to the NL820 isolator (Neurolog system, Digitimer) with gain x5. The data was digitized at 1 kHz using CED 1401 mk2 hardware and Spike2 software (Cambridge Electronic Design (CED), Cambridge, UK). The recorded data had a sampling time of 1 ms/1 kHz (Fig 1 D-E). Local field potential responses evoked by the electrical stimulation of the skin of the forepaw verified that the recording electrode was correctly placed in the forepaw area of the S1. The duration of anesthesia did not exceed eight hours. Once all the recordings were performed the animal was euthanized using a lethal dose of pentobarbital.

### Stimulation paradigm

For electrical tactile skin stimulation, pairs of intra cutaneous needles were placed at the base of the second digit of the left forepaw and at the base of the second digit of the right hind paw (Fig 2A). Single pulse stimulation was delivered to one of the stimulation sites at a time, with a pulse intensity of 0.5 mA and a pulse duration of 0.14 ms, which is above the threshold for activating tactile afferents, but well below the threshold for recruiting A-delta and C-fibers ^14,15^. A period of stimulation consisted of repeating the same stimulation at a specific frequency (0.3- 5 Hz) for 5 minutes. A stimulation period was always followed by a period of no stimulation (spontaneous activity) for about 2 minutes, and hence each stimulation period had both a preceding period and a following period of spontaneous activity against which the stimulated recording data could be compared (Fig 2B). A stimulation period of the forepaw at one frequency was followed by a stimulation period of the hind paw at the same stimulation frequency. The stimulation always started at the lower frequencies and was then gradually increased for each stimulation period for the stimulation site (Fig 2B). An evoked response to the left forepaw stimulation could be seen only in the recording electrode placed in the right S1 (Fig 1B-C). No response to the right hind paw stimulation was observed in any area, not even in the recording from the left S1, as this recording electrode was placed in the forelimb area (Fig 1B).

**Figure 2.**
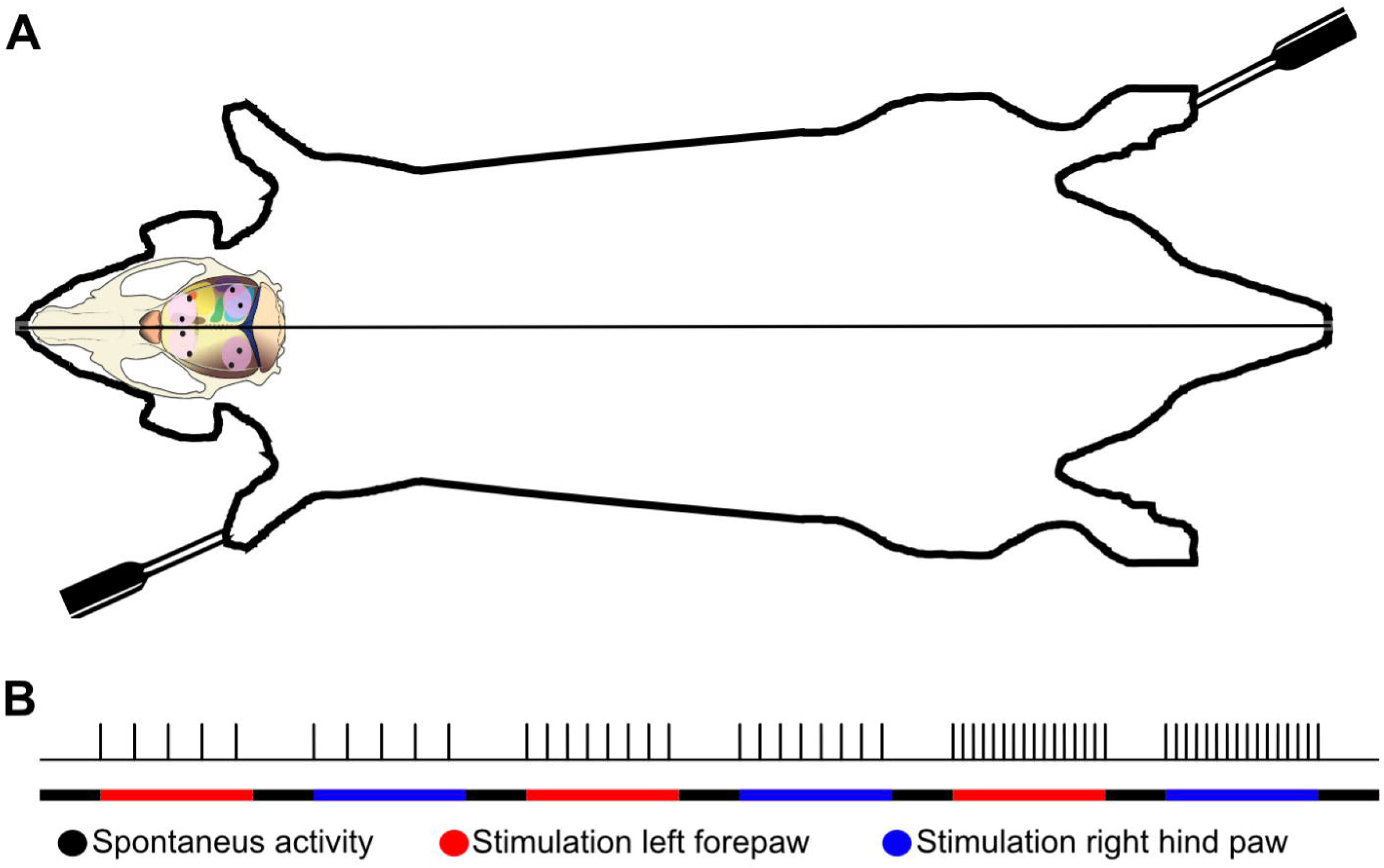
Visualization of the stimulation electrodes and the stimulation protocol. **A** Rat outline showing the placement of the stimulation electrodes in the left forepaw and right hind paw. **B** Visual representation of the stimulation protocol. The lower line shows how periods of spontaneous activity is interspersed with periods of stimulated activity and that the stimulation switched between the left forepaw and the right hind paw. As can be seen, there was a gradual increase of the stimulation frequency in consecutive periods. Each spontaneous period lasted for 2 minutes, and the stimulation periods lasted for 5 minutes. Note that for visibility, fewer impulses per period of stimulation are shown here than occurred in the actual trials. All stimulation periods were both preceded and followed by a period of spontaneous activity.

### Data collection

ECoG activity was collected from 16 of 27 animals. 11 animals were excluded due to experimental difficulties e.g., the pool leaking, electrode interference, stimulations not working properly. The stimulation followed a set protocol with alternating periods of stimulation and no stimulation as described above (Fig 2B).

### Post processing

Data was imported from Spike2 to MATLAB (MATLAB Release 2021a, The MathWorks, Inc. Natick, Massachusetts, United States) where artifacts were removed with linear interpolation between the two time-steps immediately before and after the time of external impulse. A Savitzky-Golay filter with a window size of 20 ms was used to smooth the raw ECoG data.

### Principal component analysis

Principal component analysis (PCA) was used to extract information from the ECoG channels raw data. To highlight the differences in the temporal profile we Z-scored average raw ECoG data and used an inbuilt MATLAB function (“*pca*”) to perform this analysis. The PCA was performed on the complete ECoG activity data set, both spontaneous and stimulated.

### kNN analysis

To evaluate if the cross-channel ECoG recording data points were separable between the evoked and the spontaneous activity, we performed a k-nearest neighbor (kNN) analysis on the PCA coefficients of each data point (sampled at 1 kHz), using the MATLAB classification learner toolbox, with N=5 nearest neighbors and five-fold cross-validation. The kNN analysis could be used to quantify if the activity distribution across the ECoG channels differed between the evoked and the spontaneous activity. To ensure that the results were not biased by filtering, the kNN analysis was performed on raw ECoG data. To avoid potential shock artifacts contaminating the analysis, data from the first five milliseconds after the stimulation pulse was removed from the analysis. In order to analyze the information present both in higher and lower PCs, we normalized all the time series of PC coefficients between 0 and 1, to ensure no PC was given higher or lower weight in the kNN analysis.

Stimulated activity was defined as the window size of 190 ms (5 ms – 195 ms) post stimulation pulse. The specific window size was chosen in order to avoid overlap between stimulated activity after each stimulation pulse for 5 Hz stimulation frequencies. Additionally, this window was longer than the length of any evoked responses, which lasted 20-50 milliseconds, ensuring it was not solely the presence of an evoked response that caused any separation of stimulated and spontaneous activity. As the current stimulation protocol (see “Stimulation protocol” above) comprise a larger number of spontaneous activity sample data points than the stimulated activity, we randomly chose time series within the spontaneous activity periods to obtain the same number of data points as of the stimulated activity. The kNN analysis was repeated 100 times to obtain a mean decoding accuracy for the stimulation period (i.e., how well the stimulated activity was separated from the spontaneous activity). Combined with the five-fold cross-validation, each reported kNN value represents 500 randomizations of test and training data. A new random time series of spontaneous activity was chosen every 10^th^ iteration in the classification analysis, to ensure that the accuracy reported was not a reflection of a specific subset of spontaneous activity. This analysis procedure was repeated for each individual stimulation period.

To evaluate the significance of the resulting decoding accuracy, we repeated the kNN analysis for shuffled data. A data shuffle consisted of the same data points as in the normal analysis, but with group labels (stimulated or spontaneous activity) being shuffled between them, and then we conducted a full analysis procedure for that shuffled data. The shuffling was repeated 100 times for each stimulation period and the distribution of classification rates for the shuffled data is reported in figure 5.

The kNN decoding accuracies of different stimulation periods were grouped based on the stimulation condition, and the median accuracy for all stimulation periods of that group was reported. The groups used were as follows: all stimulation conditions combined; stimulation of the left forepaw only; stimulation of the right hind paw only; and stimulation at specific frequencies, 0.3-5 Hz including data for both the forepaw and the hind paw. Wilcoxon rank-sum test was used to check the significance of the differences in accuracy observed across these stimulation conditions. Additionally, to understand the sensitivity of the kNN analysis applied to these data, we explored the effect of excluding single PCs consecutively. We also varied how many PCs that were used for the kNN. In this we started of using only the first PC and then subsequently adding each one of the lower PCs one by one, from PC 2 to PC 8. These results are reported in figure 7 and 8.

### Variations of the kNN analysis

We performed kNN analysis on the data in four different configurations, as defined by the number of compared groups and from which recording electrodes data was included.

“All areas, 2 groups”, kNN was performed on data from all eight recording electrodes and compared stimulated activity with pooled data from the two periods of spontaneous activity, that which preceded and followed the stimulation period (Fig 2B).

“All areas, 3 groups”, kNN was performed again on data from all eight recording electrodes, but now the data was compared separately to the preceding and the following spontaneous activity. This comparison was only made for stimulation frequencies below 4 Hz, due to the fact that the stimulated data, obtained during 5 minutes, was much greater than the data obtained for the spontaneous activity, which in this case lasted only for 2 minutes, for stimulation frequencies of 4 and 5 Hz.

“Non-S1”, here the kNN was performed on data from only seven of the eight ECoG recording electrodes. Data from the right S1 was excluded. In this kNN analysis, we used the pooled spontaneous activity, as in “All areas, 2 groups”.

“Only left hemisphere”, here the kNN was performed exclusively on data from the four recording electrodes placed on the left hemisphere. The kNN analysis was again performed with pooled spontaneous activity (“All areas, 2 groups”).

### Ethical considerations

All animal procedures were done in accordance with institutional guidelines and were approved in advance by the Local Ethics Committee of Lund, Sweden (permit ID M13193-2017 and M20013-2021). The animal experiments consisted of acute preparations under general anesthesia and all efforts were made to minimize suffering. All experiments were terminal.

## Results

We recorded EEG signals from eight sites across the rostrocaudal and mediolateral extent of the cortical surface (Electrocorticogram, ECoG) (Fig 1A). Weak electrical tactile stimulation of the digit skin of the left forepaw evoked distinct field potentials in the forepaw region of the right S1 (contralateral to the stimulation), but not in any other recorded area, including the left S1 (ipsilateral to the stimulation) (Fig 1B-C). Weak electrical tactile stimulation of the right hind paw did not elicit an evoked response in the left S1 (contralateral to the stimulation) nor in the right S1 (ipsilateral to the stimulation) (Fig 1B).

### PCA shows separation in covariance pattern between stimulated and spontaneous activity

For analysis of the activity distributions across the set of ECoG channels, we used time continuous ECoG recordings (Fig 1D). Episodes of spontaneous activity were mixed with episodes of tactile stimulation repeated at a specific frequency (see Table 1). Stimulations were alternately applied either to the left forepaw or to the right hind paw (Fig 2A-B). We separated the ECoG into spontaneous and evoked activity (Fig 1D-E; Fig 2B), where each stimulation period was compared to spontaneous activity that preceded and followed the stimulation period. The activity distributions across the eight ECoG electrodes were compared between these two types of activity using covariance analysis.

**Table 1.** kNN accuracy across different stimulation conditions and variations of kNN.

| Stimulation conditions | All areas, 2 groups (%) | All areas, 3 groups (%) | Non-S1 (%) | Only left hemisphere (%) | N (stimulation periods) |
| --- | --- | --- | --- | --- | --- |
| All | 72.71 | 64.65 | 66.68 | 58.84 | 102 |
| Forepaw left | 73.01 | 65.56 | 67.74 | 59.59 | 51 |
| **Hind paw right** | 72.49 | 63.27 | 66.19 | 57.15 | 50 |
| **0.3Hz** | 79.61 | 73.51 | 75.62 | 60.91 | 10 |
| **0.5Hz** | 74.53 | 62.99 | 67.96 | 57.06 | 6 |
| **1Hz** | 73.89 | 67.11 | 71.47 | 63.73 | 49 |
| **2Hz** | 67.58 | 55.13 | 61.62 | 54.12 | 8 |
| **3Hz** | 62.80 | 51.80 | 58.35 | 52.42 | 13 |
| **4Hz** | 61.29 | N/A | 57.66 | 53.34 | 8 |
| **5Hz** | 60.67 | N/A | 57.23 | 52.72 | 8 |

Median kNN accuracies for the different stimulation conditions. Column one shows the results of the kNN when the data was divided into two groups, one group of data points of the spontaneous activity and one group of data points following the tactile stimulation ('evoked' data points). Column two shows the results of the kNN when the data was divided into three groups and compared. The first group consisted of evoked data points. The second group consisted of spontaneous activity recorded just prior to the stimulation and the third group consisted of spontaneous activity in the time period following the end of the repeated tactile stimulations at the indicated frequency. Column three shows the results of the kNN when data from the ECoG electrode placed in the responding S1 was excluded (one spontaneous group). Column four shows the results from the kNN when data from the entire hemisphere contralateral to the forepaw stimulation was excluded.

The dominant covariance patterns were extracted using principal component analysis (PCA). Following PCA, each time point of the combined eight-channel ECoG recording data obtained a position in principal component (PC) space. That position in PC space could vary rapidly over time, depending on continual changes in the activity distribution across the eight ECoG channels. This caused the sequential data points to ‘wander’ in the PC space, which occurred also for the spontaneous activity but often were more pronounced following tactile stimulations (Fig 3 A-B). Whereas Figure 3 illustrates this PC space wander for individual evoked responses and its preceding spontaneous activity, Figure 4 illustrates this phenomenon for an entire stimulation period, containing 100’s of repetitions of the stimulation. Notably, the distribution of the data points during the stimulation period was different from the distribution during the spontaneous activity, even though they partially overlapped. This was the case for both stimulation of the left forepaw, which did elicit an evoked response in the right S1 recording channel (Fig 1B-C; Fig 4A), but also for the stimulation of the right hind paw, which did not elicit an evoked response in any of the recording channels (Fig 1B; Fig 4C). This separation of the stimulated versus the spontaneous activity remained also when the S1 recording channel was removed from the data set (Fig 4 B, D).

**Figure 3.**
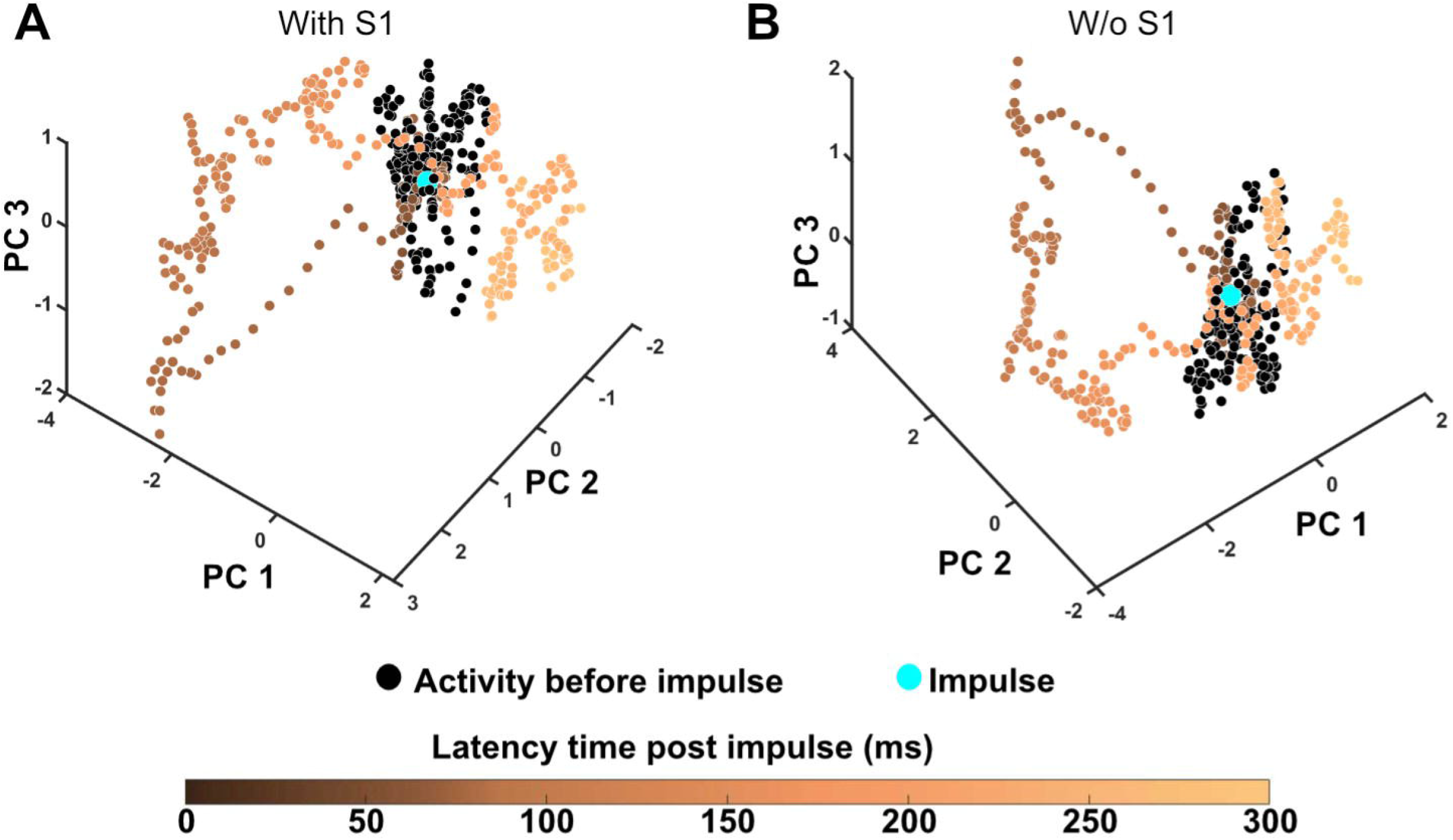
Changes in global cortical activity distribution caused by single skin stimulation impulses. **A** The change of the ECoG activity distribution, measured as the location in the PC space, evoked by a single electrical stimulation impulse to the skin of the left forepaw. The blue dot represents the time point of the stimulation. Black dots show the locations in the PC space of the data time points before stimulation, i.e., during spontaneous activity. The data points gradually change from dark (beginning) to light color (end) depending on latency time from the stimulation. Note that the post-stimulus data loops back towards the pre-stimulus data, i.e., the spontaneous activity, after about 200 ms. **B** Similar plot as in **A** but excluding the recorded data from the responding (right) S1 area. Note that the activity space occupied during the stimulation is very similar to that in **A,** but that the activity distribution now changes in a reverse cycle. In order to facilitate the comparison, the viewing angle of the plot has been somewhat adjusted from **A**.

**Figure 4.**
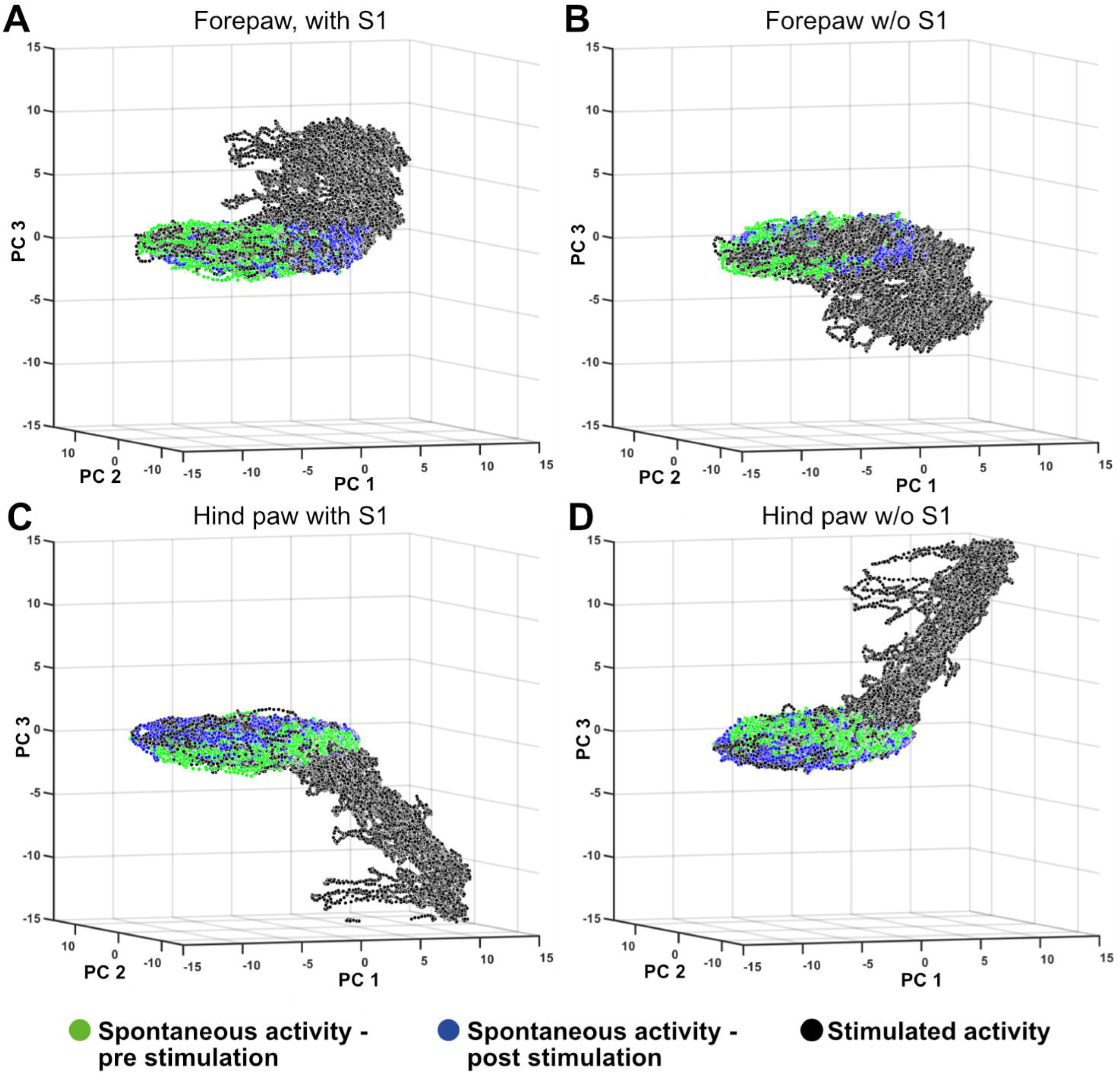
Changes in global cortical activity distributions caused by repeated single skin stimulation impulses. **A.** The change of the ECoG activity distribution, measured as the location in the PC space, evoked by electrical stimulation impulses to the skin of the left forepaw (single pulse, 0.3 Hz). The plot is similar to that in Figure 3 but shows data from the whole stimulation period (5 min) and includes both the preceding and following spontaneous activity periods. **B.** Similar 3-D plot as in **A** but omitting the ECoG recording from the right S1 area. **C, D.** Similar plots as in **A, B** but for stimulation of the right hind paw (single pulse, 3 Hz).

However, in these illustrations (Fig 3; Fig 4), we are limited by the fact that we can only visualize three dimensions simultaneously. However, the PCA could extract more covariance patterns/dimensions than these three PCs, and the natural next step for us was to quantitatively explore the maximal separation of evoked and spontaneous activity that could be obtained when also including the higher order PCs for the quantification. To this end, we applied a kNN analysis.

### Separability of ECoG covariance patterns can be evaluated using kNN

In order to quantify whether the changes in ECoG activity distributions/covariance patterns systematically differed between spontaneous and stimulated activity, we performed a k-Nearest Neighbor analysis (kNN). kNN can be used to quantify how often/what fraction of data points in neighboring parts of the PC space belong to the same category. Since we wanted to analyze whether the ECoG covariance patterns differed between spontaneous and evoked activity, the point of the kNN analysis was to quantify whether spontaneous data points were more commonly found next to other spontaneous data points rather than next to the data points that occurred during a stimulation period, and vice versa.

Whereas the kNN provided a number for the accuracy, basically the fraction of labelled data points that had a similar location (‘nearest neighbors’) as other points with the same label (spontaneous vs evoked), we also needed to compare that number with nonstructured data. To this end we used repeated shuffling of the data. For each set of shuffling we obtained a normal distribution of chance data, against which the results for the test data were compared (Fig 5A). Stimulation period data was compared against both preceding and following spontaneous periods data, either as pooled spontaneous data (‘2 groups’, Fig 5A) or as two separate groups of spontaneous data (‘3 groups’, Fig 5B).

**Figure 5.**
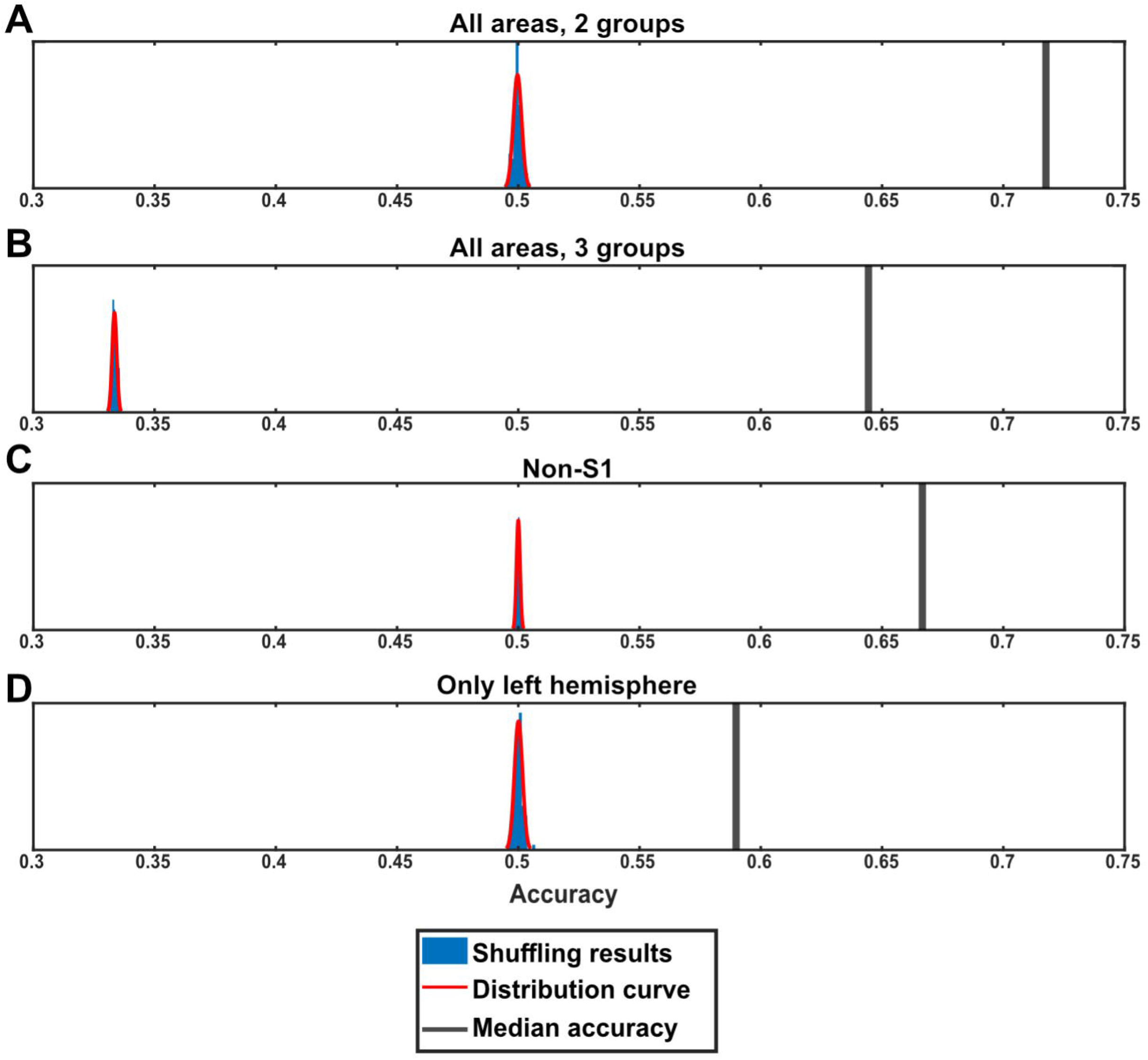
Sample kNN results for test data and shuffled data. **A.** The black line to the right indicates the median kNN accuracy for the “all areas, 2 groups”- data for the forelimb stimulation condition (all frequencies). The blue bars and the red curve indicate the kNN accuracies for the shuffled data. **B** Similar display as in A but for the “all areas, 3 groups”-data. **C** Similar display as in A but for the “non-S1”-data. **D** Similar display as in A but for the “only left hemisphere”-data.

We first looked at the combined stimulation period data from all stimulations. We found a significant median kNN accuracy when compared to the distribution of the shuffled data (Fig 5 A-D). For the “All areas, 2 groups”, the median accuracy for all stimulation periods was 72.71%, compared to the chance level of ∼50% obtained from the shuffled data (Fig 5A). For the “All areas, 3 groups” the chance level was ∼33% (0.33) and again the median accuracy across all stimulation periods was significantly higher at 64.65% (Fig 5B). Note that while the nominal median accuracy was lower for “All areas, 3 groups” than for “All areas, 2 groups”, the fact that the chance level dropped from 50% to 33% actually implies a better accuracy compared to chance for the “All areas, 3 groups” (64.65%) than for the “All areas, 2 groups” (72.71%) comparison.

Since the field potential evoked by the left forepaw stimulation in the right S1 could be prominent (Fig 1B), even though for a much shorter time (approximately 20 ms) than the duration of the time period from which the stimulated activity data points were obtained (190 ms), we also repeated the kNN analysis without including the S1 recording data (“Non-S1” kNN). Still, the ECoG covariance pattern across the included seven channels was significantly different between the evoked and spontaneous data points when comparing with the 50% chance level of the shuffled data (Fig 5C). Even when all the channels from the entire hemisphere contralateral to the forepaw stimulation were omitted, in the “Only left hemisphere” kNN, the covariance patterns between the evoked and spontaneous data were still significantly different to the chance level indicated by the shuffled data (Fig 5D).

In the account below, we consider stimulation periods with different stimulation conditions, i.e., episodes of different stimulation frequencies as well as stimulation sites (left forepaw (L-FP) and right hind paw (R-HP)) either in isolation or combined. We used Wilcoxon rank-sum test to test for significant differences in the kNN accuracy, not only for the combined stimulation conditions but also for each stimulation condition separately. Table 1 presents the median accuracies for each of the four types of kNN comparisons listed in the methods section, grouped based on stimulation conditions. These results are also visualized in Figure 6 A-D. First, stimulation periods with L-FP stimulation, which evoked a response in the contralateral S1 (Fig 1B), were compared to the accuracy of the stimulation periods with R-HP stimulation, which did not elicit a response in any recorded area (Fig 1B). We found no significant difference in the accuracy (i.e., kNN separation of the stimulated and the spontaneous activity) for these two stimulation conditions (p=0.6). When we grouped the stimulation periods based on stimulation frequency (pooling data from L-FP and R-HP stimulations), stimulation frequencies below 2 Hz had a higher accuracy than stimulation frequencies of 2 Hz and above (p<0.01) (Table. 1). This suggests that the thalamocortical system did not distribute the responses evoked at high stimulation frequencies as well as it did for lower frequencies.

**Figure 6.**
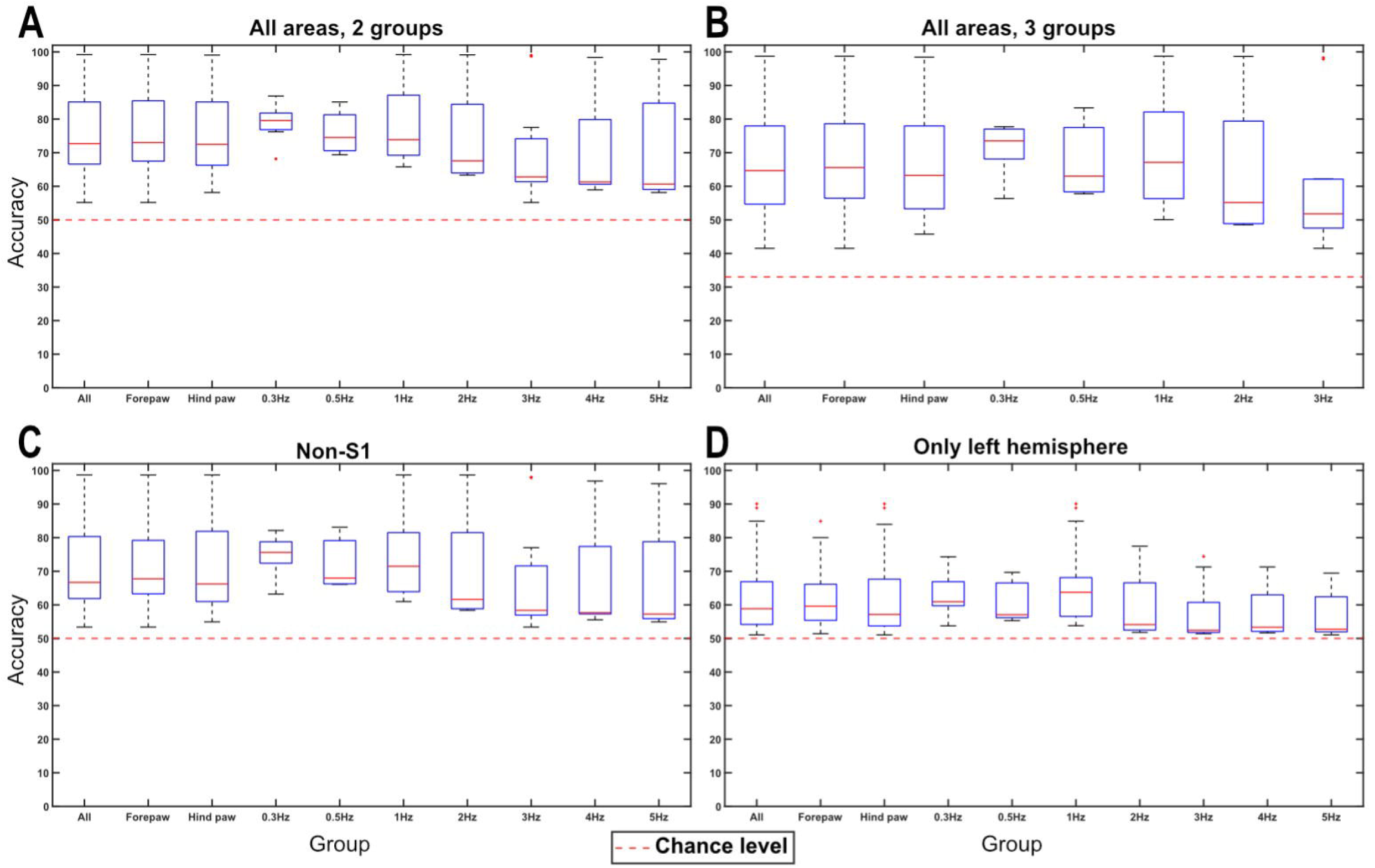
Population level kNN results. **A** kNN results from the “all areas, 2 groups”-data. The y-axis shows the accuracy (%) and the X-axis shows the stimulation conditions. For each condition, the median, quartiles, and outliers are indicated. Note that in all categories but “Forepaw” and “Hind paw”, the forepaw and hind paw stimulations are pooled, and the group contains all stimulation periods of the indicated frequency, with no differentiation between which paw the stimulation was applied to. **B** Similar display as in **A,** but for the results from the “all areas, 3 groups”-data. **C** Similar to **A,** but for the results from the “non-S1”-data. **D** Similar to **A,** but for the results from the “only left hemisphere”-data.

The Accuracy for the “Non-S1” was lower than the accuracies for “All areas, 2 groups” (Wilcoxon rank-sum test, p<0.01) (Table. 1). Similarly, “Only left hemisphere” had lower accuracies than all the other three kNNs (Wilcoxon rank-sum test, p<0.01) (Table. 1). Nevertheless, it was quite remarkable that removing the only ECoG channel showing an evoked response to either stimulation site only had a small effect on the kNN accuracy (i.e., the separation of the stimulated and the spontaneous activity). Even when we considered only the ECoG channels in the ipsilateral hemisphere, the changes in covariance patterns during the tactile stimulation were prominent. This indicates that our measure indeed did quantify the global distribution of the evoked activity.

### The kNN indicates a high dimensionality of the activity distribution

We also performed a sensitivity analysis of the kNN by removing each individual PC one at a time and explored the resulting drop in kNN accuracy. Interestingly it appeared to matter little which PC was excluded (Fig 7), each one of them had noticeable contribution to the kNN accuracy. There was no correlation between how much of the variance a certain PC explained and how much it contributed to the kNN accuracy (Pearson correlation coefficient, R<0.2). On the same theme we explored how the number of PCs used might affect the accuracy (Fig 8). This was done by at first only including the first PC and then one by one adding PC 2-8 to see how the number of PCs included affected the decoding accuracy. While increasing the number of PCs included did indeed increase the accuracy, plotting the increase in accuracy obtained for each PC added did not follow the same trajectory as the amount of variance explained by the included PC (Fig 8). While the first few PCs included caused the biggest increase in the explained variance, and this curve then started to flatten out around 5 PCs, the same first few PCs added gave the smallest increase in accuracy, and it was only around 4-5 PCs that we started seeing a larger increase in accuracy. This indicates a high dimensionality of the activity distribution, i.e., that there were multiple covariance patterns in that signal, and that a correct identification of whether the activity was evoked or spontaneous relied on information also from the higher order PCs.

**Figure 7.**
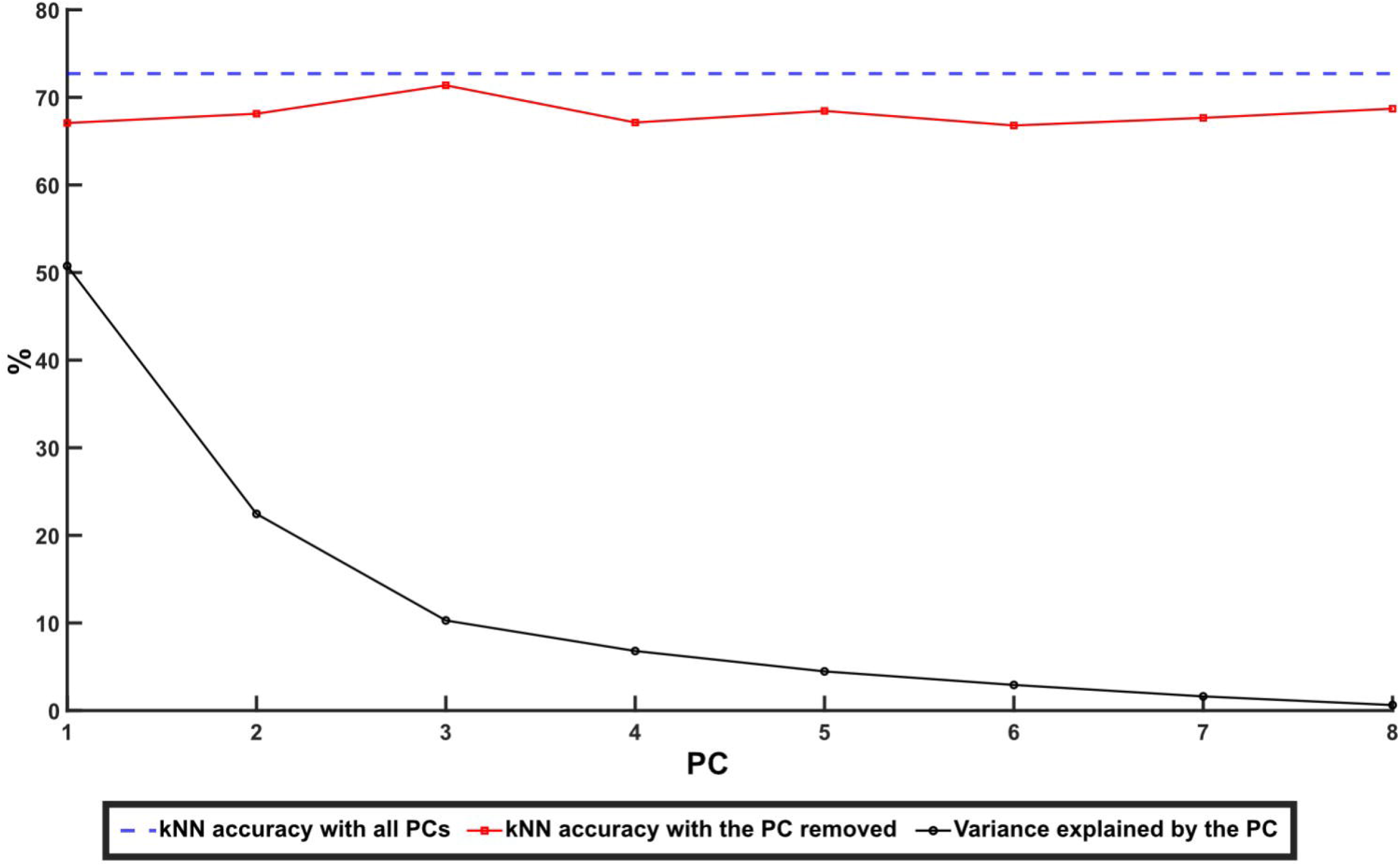
Drop in kNN median accuracy on removing specific principal components. Dashed blue line indicates the median kNN accuracy of the “all areas, 2 groups”-data with all principal components included. Red data points indicate the kNN accuracy after removing the specific PC indicated on the x-axis. Black data points indicate the variance explained by each PC.

## Discussion

We found that even weak tactile stimulations cause significant changes in the global cortical ECoG covariance patterns. Employing kNN analysis, our data was shown to have a significant separation between spontaneous and stimulated activity, also when there was no discernable local field potential response among any of the ECoG recording channels included in the analysis. These results indicate that the present method of analyzing shifts in global covariance patterns is a highly sensitive tool to identify perturbations or changes in cortical processing. Because of its low level of invasiveness (ECoG is a more highly resolved version of EEG), this is a method that could potentially also be applicable to humans, for example to study and detect perturbed cortical processing caused by neurological or psychiatric disease in EEG recordings.

### Principal component analysis and kNN

We used PCA to extract dominant patterns of cortical activity distributions (i.e., the covariance patterns across the ECoG channels) and then applied kNN analysis to investigate systematic differences in those cortical activity distribution patterns across different stimulation conditions. The kNN analysis revealed that there was a systematic shift in the activity distribution between spontaneous and stimulated activity. Notably, there was not a 100% separation between these two conditions, which suggested that the activity distribution during stimulation to some extent fell within the same patterns of activity distribution as the spontaneous activity, which we could also show (Fig 3; Fig 4). Similar findings, indicating partly overlapping activity distributions between evoked and spontaneous activity have previously been reported for cortical cell populations ^16,17^. Additionally, in these studies it was observed that the spontaneous activity featured high dimensional activity and that during sensory stimuli, the evoked activity occurred in partly orthogonal dimensions, hence increasing the dimensionality of the cell population activity ^17^. These observations are fully compatible with our results and could be the main mechanistic explanation for them. However, rather than focusing on limited cell populations, we showed that the tactile stimulation changed the activity distribution across large parts of the cortex. Thus, across-channel analysis of all eight recording channels, where each recording channel added represented an increased dimensionality in the recording data, and therefore a higher sensitivity for detecting subtle differences in cortical activity distributions, explains why the kNN accuracy consistently increased when more PCs were included (Fig 7; Fig 8).

The fact that the kNN accuracy was similar for stimulation of both the left forepaw (which evoked a field potential response in right S1) and the right hind paw (which didn’t evoke any field potential response), as well as when we removed the data from the only recording area that showed an evoked response, the right S1 (Fig 1B-C), indicated that the separation between stimulated and spontaneous activity was not due to an evoked field potential response to the stimulation in a specific recording electrode. The evoked field potential also had a much shorter duration (around 20 ms) than the time window of 190 ms included in the stimulation periods. Taken together, these observations indicate that the ECoG covariance pattern method we used here detects global shifts in the activity distribution across the cortex. Notably, the number of neurons in the neocortex, combined with the perpetually ongoing internal activity in the cortex, makes it likely that the cortical network can display a near infinite number of permutations in its activity distribution (see also Etemadi et al 2022 ^13^). The PCA will find the dominant subsets of the covariance patterns across the recording electrodes, each of which in turn represent a combination of the neuron population located under the recording electrode, and the observed separation of the spontaneous and stimulated activity patterns indicated that those covariance patterns differed between the two conditions.

The difference in kNN accuracy between different stimulation frequencies might indicate that there is a range of naturally occurring frequencies where the impact of the stimulation is greater on the network activity, thus changing the activity distribution to a greater extent. Such resonant frequencies have been shown to occur both in the thalamus and in the cortex ^18–20^. Rats under ketamine anesthesia has a prominent EEG rhythm at low frequencies, with slow oscillations peaking around 1.6 Hz ^21^, which is also where we found a higher degree of separation. It could be that stimulations in resonance with the inherent frequencies are transmitted more efficiently than other frequencies, and that this in turn results in a larger total response with a bigger impact on the global activity distribution.

### Implications for network processing

Our results are in line with the idea that the neocortex is a globally organized interconnected network ^22^ where, in principle, there is a global spread of neuronal responses to specific sensory input ^23,24^. This idea is consistent with findings that signals evoked from distant parts of the neocortex influence responses to sensory stimulation ^11,13^ and that particular aspects of evoked sensory responses can be found distributed across many parts of the cortex ^7–9,25^. Calcium imaging studies also support the notion of global processing within the cortical network ^10,17^. However, in general, such calcium imaging techniques suffer from a low time resolution, while another common method for global network analysis, EEG spectrograms, lacks the cross-channel analysis ^26^. The analysis method presented here instead allowed us to non-invasively monitor the location of the cortical network in its high-dimensional state space with a high temporal resolution. Additionally, while machine learning and artificial intelligence is becoming increasingly applied for pattern analysis of EEG data, there has been a lot of focus on preprocessing of the single channel EEG data, such as convolution or transforms, before applying the machine learning algorithm/the artificial neural network ^27^. In contrast, our analysis method only pays attention to the signal distribution data across the ECoG channels. Preprocessing could be at risk of destroying the information which is present across the EEG channels. Our method is also much less complicated than any machine learning or artificial intelligence approach would be.

In a global view of cortical computation, sensory evoked activity is continually related to and interacting with the intrinsic properties of multiple cortical subnetworks distributed globally in the cortex ^12,17^. It has been shown that spontaneous activity influences sensory responses ^28–30^. Here we have shown that it also works the other way around, i.e., even weak peripheral sensory stimulation can influence the global activity distribution compared to the spontaneous activity. Exactly what happens in the huge underlying neuronal network is of course not possible to deduce with the present non-invasive approach. Note that in comparison to the widely used spectral frequency analysis methods applied to EEG recordings, our method is ignorant of the frequency of the EEG signals. Notably, whereas EEG frequencies can be globally coordinated and therefore highly similar across channels, our analysis method instead looks at differences in the patterns of activity distribution across the channels and therefore represents information that is in principle independent of EEG spectrogram data. This also means that a general arousal effect is not what is primarily detected by this method. For general arousal to impact the patterns of activity distribution, and be detectable by our method, it would need to cause some brain areas to differentiate from other brain areas – and then it would no longer be a general arousal. The procedure to repeat the analysis for the “All areas, 2 groups” and the “All areas, 3 groups”, also controlled for, and ruled out, the possibility that the coherent period of tactile stimulation merely caused a long-term alteration in the activity distribution that persisted after the termination of a stimulation period (Fig 5B).

Additionally, when considering arousal and PCA, arousal has reportedly been correlated with the first PC of the activity distribution, which explain the majority of the variance ^17^. However, when we removed the first PC it did not result in any drastic drop in accuracy of the separation between stimulated and spontaneous activity (Fig 7). Instead, our results indicated that each individual PC alone carries little impact on the accuracy, and rather a general increase in the dimensionality of the population signal (i.e., considering many PCs in combination) is what increased the accuracy (Fig 7; Fig 8).

This leads to the question if this type of behavior would show up in the awake animal/human as well? Anesthesia would tend to reduce the general neuronal activity, as well as increasing the number of low-frequency oscillatory activity episodes associated with sleep – in this sense, the anesthesia would be expected to constrain the number of preferred activity distributions that the cortical network would spontaneously display (see also discussion in Norrlid 2021 ^12^). Hence, it could be that these effects would be somewhat harder to detect in the awake animal, though they would likely still be present. In this case, more ECoG or EEG channels may be required ^31^ for this analysis method to yield highly resolved results, as increased dimensionality increased the sensitivity (Fig 8). This increase in sensitivity is also why this method could work in EEG as well as in ECoG, despite EEG being noisier and more low resolving than ECoG, as the higher number of recording channels in standard EEG approaches would increase the dimensionality and thus sensitivity (Fig 8) of the method. In the awake human, where there may be a more reliable conscious control of the initial internal cortical activity in relation to an (anticipated) sensory input, it could instead be possible that the effects we observe here would be easier to detect, and thus require fewer EEG channels.

**Figure 8.**
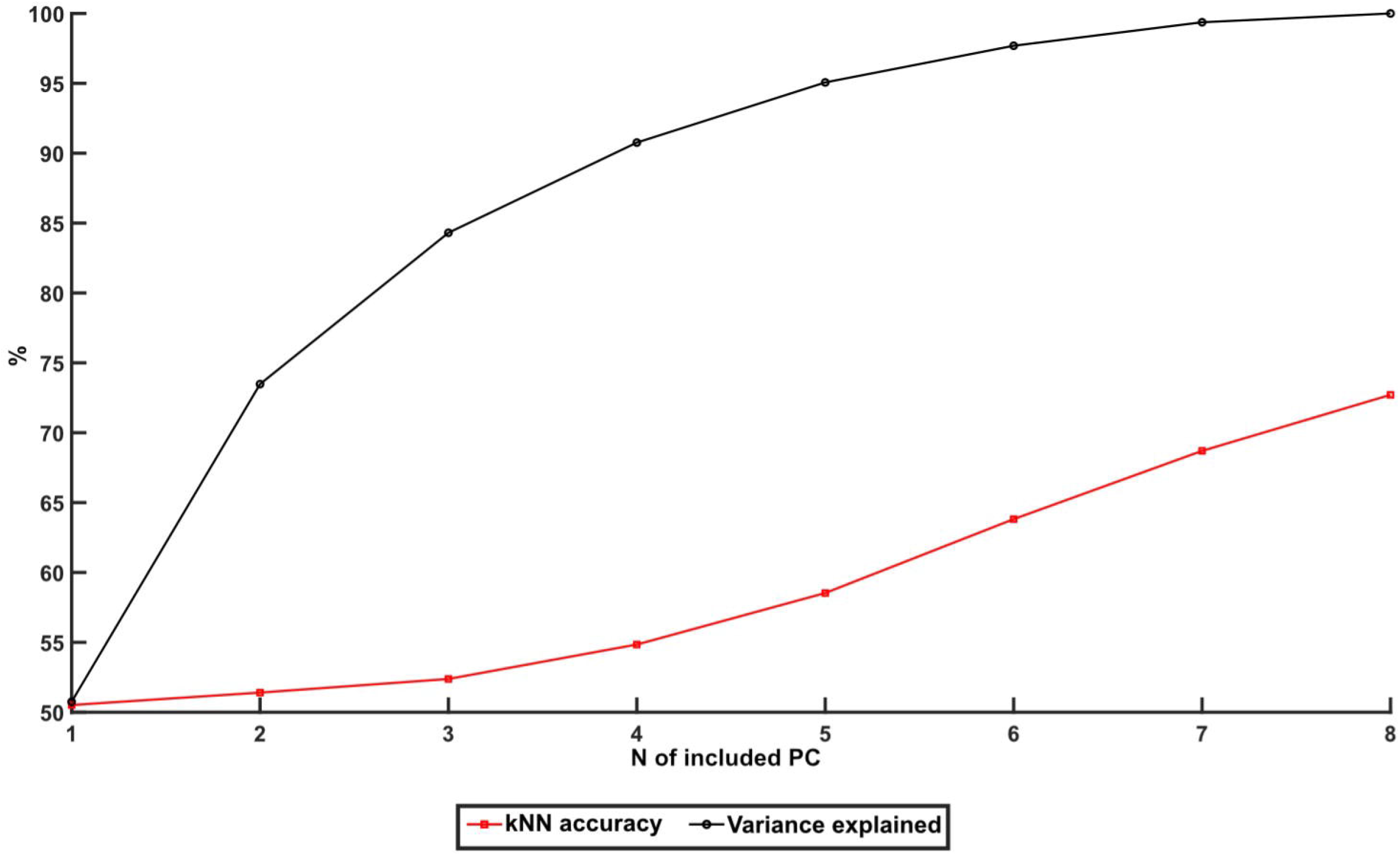
kNN accuracy per each added principal component. Median kNN accuracy for the “all areas, 2 groups”-data as a function of the number of PCs included in the kNN analysis (red data points). The black data points illustrate the variance explained as a function pf the number of PCs included.

### Concluding remarks

In conclusion, our results further strengthen the theory that the entire neocortex can be seen as one globally organized interconnected network. In addition, the fact that we could record global changes in activity distribution to a weak tactile input using non-invasive techniques suggests that it should be possible to do this also with skull surface EEG recording methods, opening up for applications for detection of even small activity changes, which in turn could be relevant for monitoring a number of neurological and psychiatric disease states.

## Limitations of the study

For practical considerations, such as spatial constraints within craniotomies and the available space above the skull, as well as the inherent limitations of current recording equipment, our investigation was constrained to the utilization of eight recording electrodes, thereby permitting the simultaneous recording of neural activity from eight discrete anatomical regions. In the pursuit of future research endeavors, it would be of interest to investigate if the quantity of electrocorticography (ECoG) channels, recording electrodes, could be increased.

Moreover, it was necessary to employ anesthesia in our study. This could potentially limit the study as the range of EEG-activity is limited to sleeping patterns. A natural next step would be to employ the same type of analysis on EEG-recordings from awake humans.

## Acknowledgments

This work was funded by the Swedish Research Council VR, project no. 2019-01623. We thank Dr. Anders Wahlbom for help with the experimental set-up and initial data handling.

## Author contributions

Conceptualization H.J. and F.B.; Methodology A.M., U.R., H.J. and F.B.; Software A.M. and U.R.; Validation A.M.; Formal Analysis A.M.; Investigation A.M. and F.B.; Resources H.J. and F.B.; Writing – Original Draft A.M. and F.B.; Writing – Review & Editing A.M., U.R.,

H.J. and F.B.; Visualization A.M.; Supervision F.B.; Funding Acquisition H.J. and F.B..

